# Protein Charge Neutralization is the Proximate Driver Dynamically Tuning a Nanoscale Bragg Reflector

**DOI:** 10.1101/2021.04.23.441158

**Authors:** Robert Levenson, Brandon Malady, Tyler Lee, Yahya Al Sabeh, Phillip Kohl, Youli Li, Daniel E. Morse

## Abstract

Reflectin is a cationic, block copolymeric protein that mediates the dynamic fine-tuning of color and brightness of light reflected from nanostructured Bragg reflectors in iridocyte skin cells of squids. In vivo, neuronally activated phosphorylation of reflectin triggers its assembly, driving osmotic dehydration of the membrane-bounded Bragg lamellae containing the protein to simultaneously shrink the lamellar thickness and spacing while increasing its refractive index contrast, thus tuning the wavelength and increasing the brightness of reflectance. In vitro, we show that reduction in repulsive net charge of the purified, recombinant reflectin – either (for the first time) by generalized anionic screening with salt, or by pH titration - drives a finely tuned, precisely calibrated increase in size of the resulting multimeric assemblies. The calculated effects of phosphorylation in vivo are consistent with these effects observed in vitro. X-ray scattering analyses confirm the sphericity, size and low polydispersity of the assemblies. Precise proportionality between assembly size and charge-neutralization is enabled by the demonstrated rapid dynamic arrest of multimer growth. The resulting stability of reflectin assemblies with time ensures reciprocally precise control of the particle number concentration, thereby encoding a precise calibration between the extent of neuronal signaling, osmotic pressure, and the resulting optical changes. The results presented here strongly suggest that it is charge neutralization, rather than any change in aromatic content, that is the proximate driver of assembly, fine-tuning a colligative property-based nanostructured biological machine. A physical mechanism is proposed.

## Introduction

Color and brightness of light reflected from skin cells of the shallow-water Loliginid squids are dynamically tunable for adaptive camouflage and communication^[1]^. Prior research showed that acetyl choline (ACh), released from neurons emanating from the brain to enervate patches of skin cells, activates muscarinic ACh receptors, triggering a signal-transduction cascade that culminates in phosphorylation of the reflectins, the major constituent of the intracellular, membrane-enclosed lamellae of nanoscale Bragg reflectors in the iridocytes^[2–4]^. The resulting assembly of the reflectins triggers the efflux of water from their membrane compartments, simultaneously shrinking the thickness and spacing of the Bragg lamellae while increasing their refractive index contrast, increasing the brightness of reflectance while tuning its color across the visible spectrum, from red to blue^[4,5]^. These effects proved to be fully reversible and repeatedly cyclable. These processes also were found to drive a corresponding dehydration of the multiple reflectin-containing vesicles within the tunable leucophores (found thus far only in the females of one species), with the heterogeneously sized pycnotic vesicles acting as bright, white, broad-band Mie reflectors^[6]^. Reflectin thus is seen to act as a signal-responsive molecular machine that regulates an osmotic motor to tune nano-photonic reflectors.

Multiple mechanisms have been implicated^[7–11]^, but uncertainty remains about the proximate trigger that first drives assembly. Xie and colleagues showed that small aromatic molecules including imidazole can drive assembly^[7]^, and suggested a similar process might initiate assembly in the physiological control of photonic behavior[7]. In contrast, Izumi *et al.* and DeMartini *et al.* observed that neuronal triggering of the reflectin-mediated photonic changes requires ACh-induced site-specific phosphorylation of the reflectins, and proposed that this phosphorylation overcomes Coulombic repulsion of reflectin’s observed excess positive charges to permit condensation and assembly^[2,4]^. Levenson *et al.* then showed that progressive reduction of reflectin’s positive charge – either by pH titration or by genetic engineering - drives proportional assembly of the initially disordered protein^[8,11]^. However, because titration reduced the charge of the imidazolium ions of histidine residues, and the genetic engineering secondarily affected the net balance of aromatic residues, some questions still remained. Accordingly, we chose to investigate the effects of charge-neutralization of reflectin in greater depth.

## Results and Discussion

### Titration is a surrogate for phosphorylation, driving proportional assembly

As shown here by dynamic light scattering (DLS) (**Figure 1A**), and previously demonstrated^[8,11]^, pH-titration of reflectin’s excess positive charges serves as an *in vitro* surrogate for the neutralization by phosphorylation in vivo, driving progressive assembly of the purified protein with increasing pH. For the experiments shown here, purified *D. opalescens* reflectin A1 was first extensively equilibrated by dialysis into 25 mM sodium acetate buffer at pH 4.5; reflectin remains stably monomeric under these acidic conditions. Assemblies formed by pH neutralization are essentially stable with time. Comparison of results from three buffer systems demonstrates relatively little buffer-specific effects. Using the known pK_a_’s of the constituent amino acid residues allows us to calculate the net charge density (net positive charge/100 residues) of the protein for each measured value of pH^[8,11]^, permitting comparison of the observed effects of neutralization by titration with those that might be expected as a result of the in vivo phosphorylation with 1, 2, or 3 phosphates per reflectin (as quantitatively measured by 2D electrophoresis and mapped by LC-MS^[2,12]^) (**Figure 1B**). These results reveal a quantitatively predictive relationship between net charge density and reflectin assembly size over the range predicted for the physiological phosphorylation. They also confirm the quantitative relationship between charge and size revealed in our analyses of phosphomimetic, deletion, and other mutants^[8,11]^.

**FIGURE 1.**
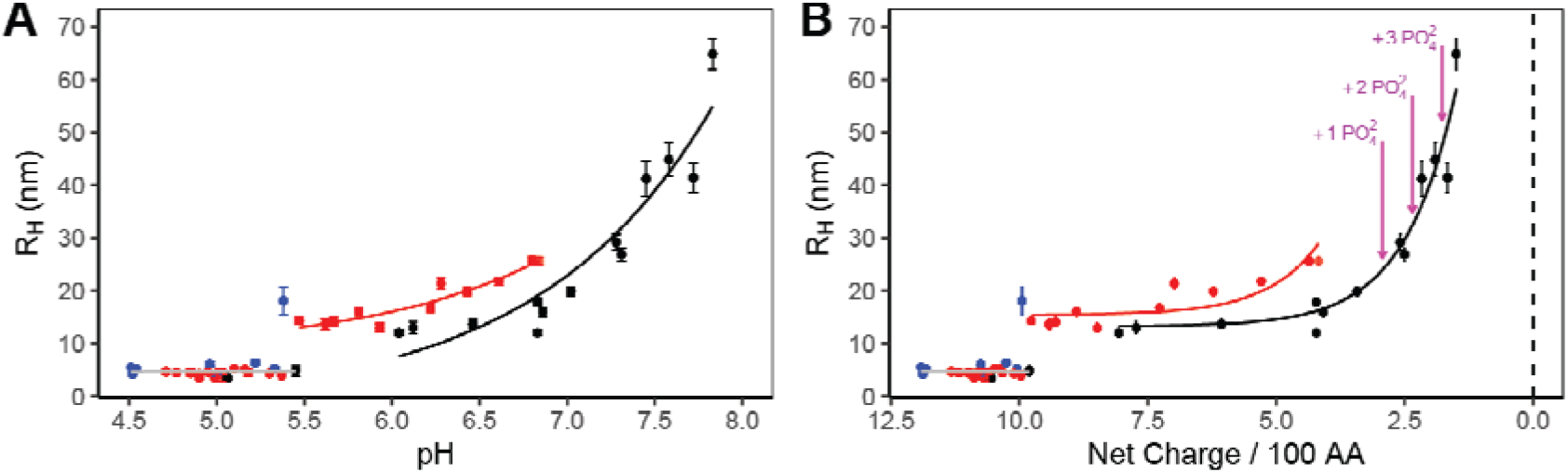
**A)** Assembly sizes, measured by DLS, of *D. opalescens* reflectin A1 upon dilution into 25 mM buffers (Blue = sodium acetate; red = MES; Black = MOPS). A1 was dialyzed into 25 mM sodium acetate, pH 4.5, before pH neutralization to labelled pH conditions. Each data point corresponds to the mean of 10+ minutes of continuous DLS measurements of a freshly pH-neutralized sample. Error bars show the standard deviation of measurements. **B)** Data from panel A plotted as a function of calculated protein linear net charge density (Details in Methods). Calculated net charge density of reflectin A1 at various stages of physiological phosphorylation (at pH 7) indicated with purple arrows.

### Effective neutralization by anionic screening drives assembly

Assembly of reflectin also is driven by anionic screening of its positive charges, resulting in an effective neutralization of electrostatic repulsion that is similar to, and interacts with, the effect of pH-titration (**Figure 2**). For these experiments, purified reflectin A1 was first extensively equilibrated by dialysis into 25 mM sodium acetate buffers at pH 4.0, 4.5, and 5.0. Dilution into the corresponding buffers containing varying concentrations of NaCl then triggered the appearance of turbidity (measured as absorbance at 350 nm) at pH-dependent salt concentration thresholds, indicating assembly, with salt concentration thresholds decreasing as pH is increased (**Figure 2A**).

**FIGURE 2.**
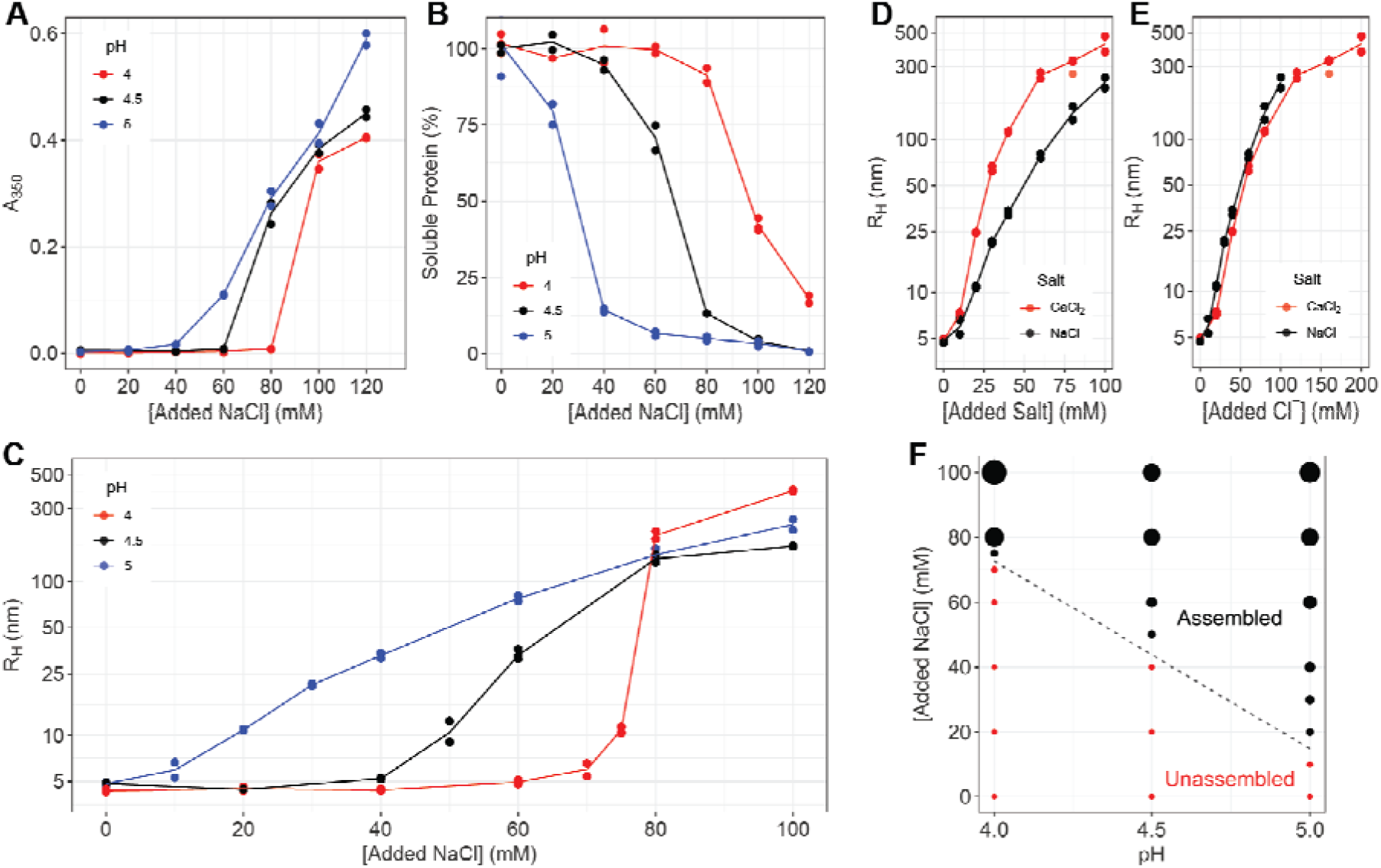
**A)** Turbidity (A_350_) of reflectin A1 solutions in 25 mM sodium acetate as a function of pH and NaCl concentration. **B)** Percent of soluble protein remaining in the supernatant after centrifugation of samples in A, relative to parallel controls with no salt. **C)** Sizes of majority populations of assembled reflectin assembl es, as measured by DLS. Each data point is an average of 40 min continuous DLS measurement (see Methods). **D-E)** Comparison of NaCl- and CaCl_2_-induced assembly at pH 5.0, as a function of **D)** salt concentration and **E)** Cl^-^ anion concentration. **F)** Reflectin A1 assembly as a function of pH and NaCl concentration. Point sizes are scaled to particle size as measured by DLS. For this panel, all particles with R_H_ > 7.5 nm are considered assembled.

Centrifugation analyses confirmed that turbidity was due to the formation of precipitable protein assemblies (**Figure 2B**). Virtually all of the reflectin was removed by centrifugation of precipitate at higher salt concentrations and pH, demonstrating assembly of the bulk population in accord with the turbidimetric data. Measurement of these samples by DLS confirmed assembly above the threshold salt concentrations, showing the size of these reflectin assemblies increasing reproducibly and progressively with salt concentration (**Figure 2C**). These salt-driven assemblies of tunable size consisted of single majority populations as judged by DLS, with minority populations of larger particle sizes occasionally observed at lower salt concentrations. The majority populations were stable over ≥40 min (not shown), suggesting that they undergo dynamic arrest similar to that undergone by assemblies formed through pH-neutralization (cf. below). Below the salt concentration thresholds for assembly, DLS shows a majority population of stable particles of R_H_ = 4-5 nm, indicating the stability of the unstructured monomers under these conditions. In contrast, parallel analyses of bovine serum albumin (BSA; not shown) under identical buffer conditions showed no aggregation or assembly at all values of pH and salt concentration tested.

Significantly, twice as much monovalent NaCl is required to drive the same extent of reflectin assembly as driven by divalent CaCl_2_ (**Figures 2D, E**), showing unequivocally that it the increasing concentration of the anion (Cl^-^) that is driving assembly, by progressively screening (effectively neutralizing) the effect of reflectin’s excess positive charges.

Plotting of assembly size as a function of both pH and NaCl concentration shows that the effects of charge screening and pH are interdependent, with both the salt concentration threshold and range for tunable assembly varying systematically with pH (**Figure 2F**). The progressively lower threshold concentration for assembly at progressively higher pH is readily explained by the progressive neutralization of the protein with increasing pH, thus reducing the requirement for neutralization by charge-screening. TEM analyses (not shown) of negatively stained reflectin A1 at pH 5.0 confirm the formation of assemblies with spherical morphologies exhibiting progressively larger sizes with progressively higher salt concentrations, consistent with assemblies formed by pH-neutralized reflectin A1 wildtype and mutants^[8,11]^.

### X-ray scattering confirms results of dynamic light scattering

Synchrotron small angle X-ray scattering (SAXS) data collected from reflectin samples in a pH-neutralized, multimeric state show characteristic features of sphere-like particles (**Figure 3**). The observed set of broad peaks in the scattering data can be attributed to form factor scattering arising from the Fourier Transform of a solid sphere. The data fit closely to a model (solid line) consisting of a group of randomly dispersed solid spheres, with an average radius = 23.2 ± 3.0 nm. The fit of the data to this model is remarkably faithful except at very low Q range, where deviation can be explained by the presence of a small number of larger aggregates that were not included in the model. The average radius measured by SAXS agrees well with the R_H_ (20 nm) measured by DLS for the same sample, and the observed polydispersity (ca. 13%) is consistent with previous estimates by DLS and TEM^[11]^. Samples prepared under different conditions exhibited a similar low polydispersity, with calculated spherical radii also closely consistent with the R_H_ values measured by DLS.

**FIGURE 3.**
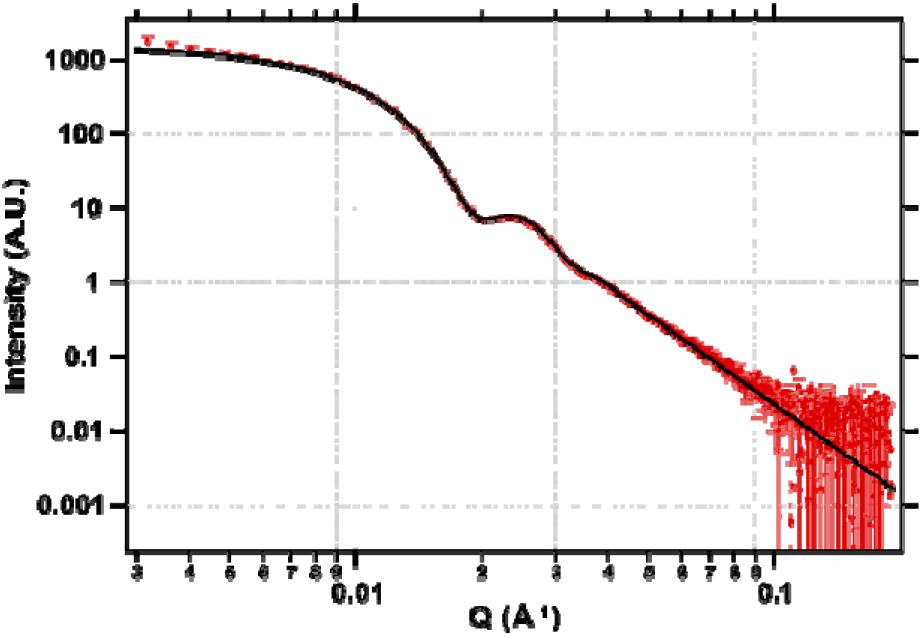
Synchrotron SAXS data for reflectin A1 multimers (4 mg/mL) in 15 mM MOPS, pH 7.5, after subtraction of buffer background. The data were fit to a spheroid model using SAXS modelling package IRENA, yielding an average particle radius of 23.2 nm + s.d. = 3.0 nm. DLS indicates the same sample to have R_H_ = 20 nm.

### Reflectin assemblies are stabilized by dynamic arrest

Data shown in **Figure 4** show the rapid dynamic arrest of growth and stabilization of reflectin assemblies. Incremental additions of reflectin monomer added successively to a buffered solution at pH 7.5 do not progressively augment the size of the initially formed multimers, but instead assemble independently upon each new addition, forming new assemblies of approximately constant size as measured by DLS (**Figure 4A**). The total particle scattering signal, a function of both particle size and concentration, increases linearly with new incremental additions of protein, showing that each newly added aliquot of reflectin does not substantially aggregate with the previously added population or precipitate (**Figure 4B**). This also is seen by comparison of changes in the intensity and volume distributions as a function of the incremental additions (**Figure 4C-D**). Significantly, the constancy of sizes indicated in the volume distribution (**Figure 4D**) confirms that the majority of reflectin monomers independently assemble to the same predetermined size upon each new addition to the same solution at fixed pH. Features of the reflectin sequence predisposing it to rapidly form a network of extensive, non-covalent cross-links that may contribute to such dynamic arrest are considered in the Discussion.

**FIGURE 4.**
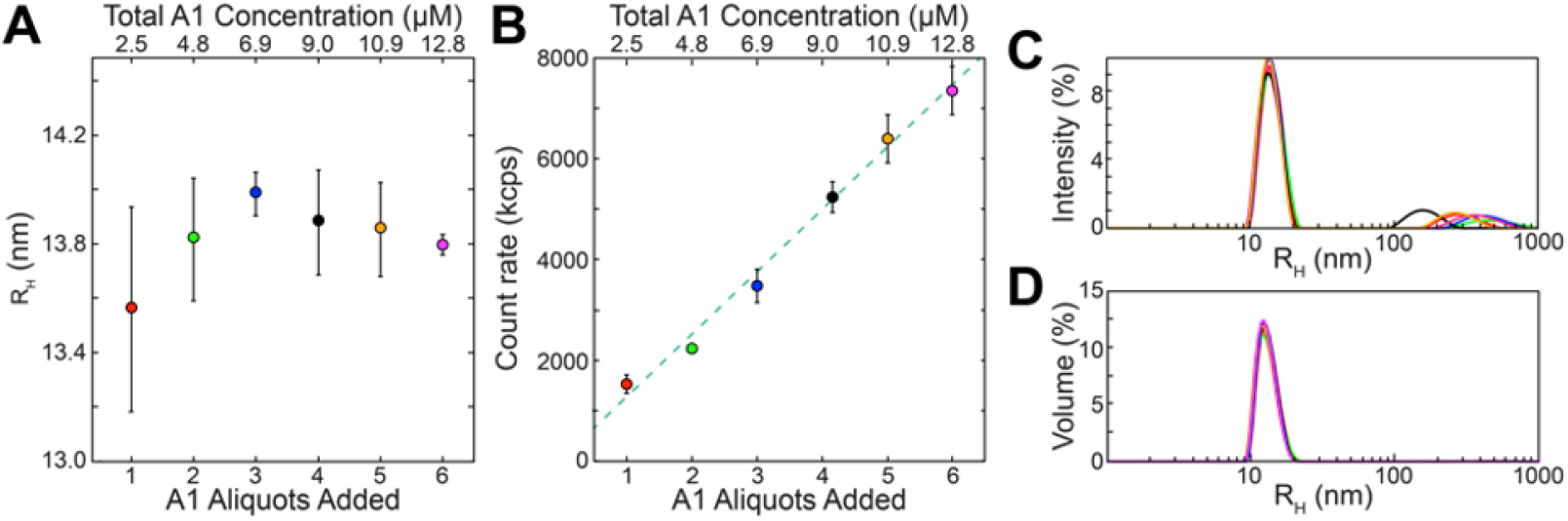
Assemblies of reflectin are dynamically arrested and stable, as seen in these results of incremental additions of reflectin A1 monomers to the same 5 mM MOPS, pH 7.5 (neutralizing, assembly-driving) solution. **A)** Reflectin assembly sizes (measured as the predominant DLS volume distributions; cf. Methods) as a function of monomer aliquots added. **B)** Total scattering count rate as a function of monomer aliquots added (bottom x axis) and cumulative concentration (top × axis). Each point in A-B is the average of 3 replicate experiments, in each of which every aliquot addition was analyzed by 3 individual DLS measurements; error bars signify ± one S.D. between averages of replicates. Samples showed no significant variation over time following each aliquot addition, consistent with previous work^[8,11]^. Representative Intensity **(C)** and Volume **(D)** distributions observed after addition of the 6th aliquot of monomer, with results after each aliquot shown in a different color.

## Discussion

The data presented here show that charge screening by added salt anions – neutralizing the effect of charge while leaving the imidazolium ions intact – drives assembly of reflectin in a manner closely comparable to the previously reported effects of pH titration and genetic engineering^8,11^, and that the effects of salt concentration and pH are interdependent. Analyses of assembly driven by pH titration (**Figure 1**) show comparable results with three buffer systems, ruling out any major effects of the buffers used. (The small discontinuity at ca. pH 5.5, which is observed consistently^[11]^, is formally indicative of an energy barrier to initial assembly.) Analyses in terms of reflectin’s effective repulsive net charge densities confirm the previously reported, predictive and precisely proportional relationship between the extent of net charge density neutralization and the size of the resulting reflectin assemblies, and demonstrate that these observations are consistent with those predicted for the physiologically observed phosphorylation^[2,11]^ of reflectin. X-ray scattering confirms the sphericity, size and relatively low polydispersity of the assemblies previously determined by DLS and TEM measurements^[8,11]^. Considering that the x-ray beam measures the ensemble average of the entire illuminated volume (~0.2 mm × 0.2 mm × 1.0 mm) of the liquid sample containing trillions of particles, the relatively low size dispersion (± 13%) indicates the operation of a highly effective, global mechanism for size control of the multimeric state.

Our results demonstrate unequivocally that charge neutralization of reflectin A1 is the proximate trigger of assembly, and that the extent of charge neutralization proportionally controls the size of the resulting assembly. This precise relationship depends upon the rapid dynamic arrest and subsequent stability of assembly size. At low pH and low salt concentrations, condensation, folding and assembly all are inhibited by electrostatic repulsion between cationic side chains, present in high proportion in the reflectins^[8,9,13]^. Reduction of this repulsion by phosphorylation *in vivo,* or by pH-titration, charge-screening (or genetic engineering^[11]^) *in vitro,* allows attractive chain interactions to quickly drive condensation, folding and assembly, with rapid arrest by a network of interactions that inhibits further dynamics and growth. Formally, this dynamic arrest of assembly is determined by the balance of short-range (weak) attractive forces and long-range (strong) repulsive forces, as well understood for many colloidal systems^[14]^. In the case of reflectin assembly, these weak attractive forces are likely to include a combination of hydrogen- and hydrophobic bonding, β-stacking, coil-coil interactions, cation-π sulfur-π π-π van der Waals and other forms of noncovalent bonding, while coulombic repulsion likely comprises the dominant strong interaction^[9,11]^.

This precise relationship between effective charge and size has a profound, direct bearing on the physiological mechanism by which reflectin controls biophotonic behavior: Because the size of the reflectin multimers is directly and inversely proportional to the *number-concentration of reflectin particles* within their membrane-bounded compartment, the charge vs. size relationship directly controls the osmotic pressure within that compartment, and thus precisely controls the osmotic dehydration of that compartment - driving the observed changes in the wavelength and intensity of the reflected light. The resulting dependence of this colligative property – osmotic pressure – on the extent of phosphorylation of reflectin thus ensures a precisely calibrated relationship between the neuronal delivery of ACh and the resulting color and brightness of the iridocyte. [Two other mechanisms contribute to the signal-dependent dehydration of the Bragg lamellae: “dewatering” of the reflectin molecules by steric exclusion resulting from the induced folding and hierarchical assembly of the protein, and the Gibbs-Donnan re-equilibration resulting from the similar steric displacement of some of the protein’s neutralizing small counterions, transmembrane migration of these ions to maintain electrical neutrality, and the consequent movement of water to equalize osmotic pressure[4]. Both mechanisms would operate in strict proportionality to the extent of assembly as well.]

We previously discussed the detailed mechanisms controlling reflectin’s assembly after activation by charge-neutralization, with data indicating that neutralization of the block copolymeric protein’s cationic, coulombic repulsion permits sequential condensation and folding, with the emergence of previously cryptic hydrophobic surfaces that are likely to facilitate hierarchical assembly^[2,8,9]^. The block copolymeric structure of reflectin A1 can thus be thought of as a concatenation of alternating, opposing expansion and contraction springs: coulombic repulsion of the cationic linkers (expansion) keeps the molecule in an extended and intrinsically disordered state until charge neutralization sufficiently opposes that repulsion, relaxing the stress on the conserved domains to allow the entropic drive encoded in their sequences to trigger condensation and secondary folding (contraction), causing emergence of hydrophobic surfaces and/or beta structures that facilitate hierarchical assembly^[8,9]^.

Beyond their physiological roles in biophotonics, reflectin proteins have been shown to form materials of various structures including optical gratings, fibers, and thin films, all with properties responsive to their environment^[15–20]^. Additionally, they’ve been shown to serve as substrates for cell growth^[21]^ and to act as proton conductors and transistors^[22–24]^.

It is interesting to note that the repeated anionic domains in an intrinsically disordered melanosomal protein recently have been found to act as the charge sensors controlling acid-triggered amyloidogenic assembly^[25]^, analogous to the role of the repeated cationic linkers that act as the charge sensors controlling reflectin’s assembly^[11]^. While numerous examples of pH- and charge-regulated transitions of protein assembly states are known^[26–30]^, and this principle recently was used to design synthetic peptides with multiple histidines buried with hydrogen-bonded networks to realize preprogrammed, pH-driven conformational changes^[31]^, this is the first report of which we are aware of a genetically preprogrammed, precisely controlled dependence of the size of protein assembly upon net charge density, with direct connection to the mechanism of control of biological function *in vivo.*

## Conclusion

Charge neutralization by phosphorylation triggers a precisely limited assembly of the cationic, block-copolymeric protein, reflectin, to drive a calibrated osmotic tuning of color reflected from Bragg lamellae in cells of squid skin^[2–6,8,11]^. Dynamic light scattering analyses show here that effective neutralization by anionic screening provides a surrogate for phosphorylation, driving reflectin’s proportional assembly *in vitro*. Because anionic screening is sufficient to drive reflectin assembly with no change in the protein’s histidine imidazolium ions or other aromatic residues, we conclude that charge neutralization and its consequent neutralization of Coulombic repulsion alone are the proximate driver of assembly, with no requirement for additional aromatic compounds as suggested by Xie *et al.*^[7]^. Assembly driven by effective neutralization by charge screening and by pH-titration of the protein’s imidazolium ions of histidine residues are interdependent, as expected, and the calculated effects of phosphorylation *in vivo* are shown to be consistent with these effects observed *in vitro*. Small-angle X-ray scattering confirms the sphericity, size, and low polydispersity of the assemblies. Precise proportionality between the extent of charge-neutralization and assembly size is shown to result from rapid dynamic arrest of assembly and subsequent stability, ensuring precise calibration between the initiating neuronal signal triggering reflectin phosphorylation^[2,6]^ and the resulting, osmotically fine-tuned color reflected from the reflectin-containing Bragg reflector *in vivo*. A physical mechanism for the continually fine-tuned control of reflectin’s assembly by charge, considering the protein’s alternating block-copolymeric domains acting analogously to a concatenation of alternately opposing springs, is discussed

## Materials and Methods

### Reflectin expression, purification, and sample preparation

Recombinant reflectin A1 wild-type from *Doryteuthis opalescens*, with no affinity tags, was expressed in *Escherichia coli*, resolubilized from inclusion bodies, chromatographically purified and lyophilized as described previously,^[8]^. Electrospray ionization mass spectrometry confirmed expression of the full-length protein, with purity verified by SDS-PAGE as previously^[8,11]^.

For analyses, lyophilized purified reflectin was solubilized in either 0.22 μm-filtered 25 mM acetate, pH 4.5, or 0.22 μm-filtered H_2_O (below). Concentration was determined spectrophotometrically from A_280_ using a calculated extinction coefficient of 120,685 M^−1^ cm^−1^. Protein was stored at 4 °C between use. Buffers and water were all filtered (0.22 μ m), centrifuged (10 min at 18,000 × g) and equilibrated to room temperature before use.

For pH-driven assembly using sodium acetate, MES, and MOPS buffer, and for salt-driven assembly, purified reflectin was first solubilized in 25 mM sodium acetate, pH 4.5. The sample was then equilibrated by 3 successive dialyses (1:1000) against the same buffer; and then either used directly or dialyzed 3 times (1:1,000) into 25 mM sodium acetate buffer at pH 4.0 or 5.0. DLS verified that no assembly or precipitation occurred during these buffer exchanges. Reflectin concentration was determined spectrophotometrically as above, diluted to 100 μm, and this stock solution was then diluted 10-fold to 10 μM into either a 25 mM buffer solution of higher pH, or a pre-equilibrated sodium acetate and salt solutions of same pH. For pH-neutralized samples, solution pH was measured immediately after DLS analysis using a calibrated micro pH electrode (Mettler Toledo InLab Micro). BSA controls (using pure lyophilized BSA from Thermo Fisher) were identically prepared in 25 mM sodium acetate, pH 4.5.

For SAXS analyses, lyophilized reflectin was solubilized in 0.22 μm-filtered H_2_O, adjusted to 200 μM, and diluted to a final concentration of 4 mg/mL (90 μM) in 15 mM MOPS, pH 7.5.

To evaluate the effects of successive additions of reflectin at a fixed pH, 1.25 μL aliquots of 81 μM H_2_O-solubilized reflectin were sequentially added to 40 μL mM MOPS pH 7.5 in the DLS cuvette. Size after each addition was measured continually for ≥ 10 min, after which the next aliquot was added and measured. Total scattering count rates were obtained from the DLS instrument software.

### Dynamic light scattering

DLS was performed using a Malvern Zetasizer Nano ZS (Worcestershire, UK). Samples (40 μL) were measured continuously at 25 °C, beginning immediately after protein addition, to ensure equilibration and stability.

### X-ray Scattering

Synchrotron Small-Angle X-ray Scattering (SAXS) was analyzed at the Stanford Synchrotron Radiation Lightsource. Data reduction and model fitting were carried out using the NIKA and IRENA software packages from Argonne National Laboratory^[32,33]^.

### Turbidity and Centrifugation

Solubilized and equilibrated protein was diluted into 25 mM sodium acetate ± salt to yield 40 μL of 10 μM reflectin. After 10-min at 20 °C, A_350_ was measured with a Thermo Fisher Nanodrop spectrophotometer. Samples then were centrifuged 10 min at 18,000 × *g* and the supernatant A_280_ measured relative to a parallel control in buffer with no added salt.

## Acknowledgements

We thank Mike Gordon, Roger Hanlon, and Steve Senft for their helpful discussions that improved our manuscript. This research was supported by the U.S. Army Research Office under cooperative agreement W911NF-19-2-0026 for the Institute for Collaborative Biotechnologies, and by grants from the U.S. Army Research Office (W911NF-17-1-0160 and W911NF-20-1-0257) and the U.S. Dept. of Energy, Basic Energy Sciences (DE‐SC0015472). Use of UCSB’s Materials Research Laboratory Shared Experimental Facilities was supported by the Materials Research Science and Engineering Center Program of the National Science Foundation under Materials Research Award 1121053.

## Table of Contents Figure

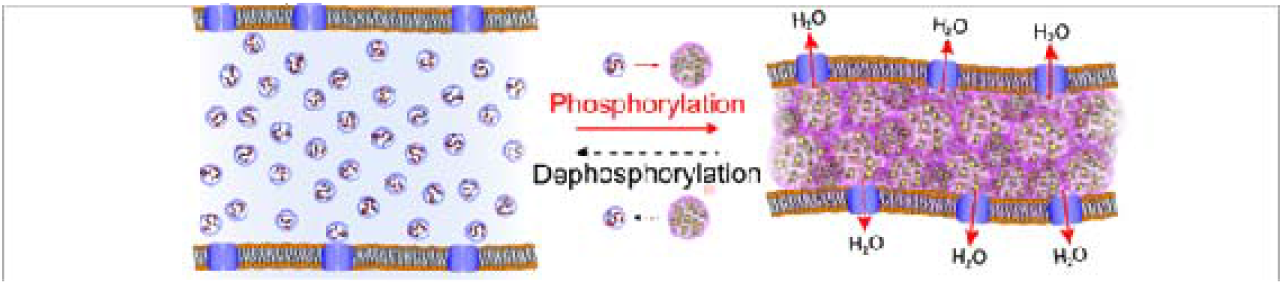

Charge neutralization by phosphorylation triggers a precisely limited assembly of the cationic, block-copolymeric protein, reflectin, driving a calibrated osmotic tuning of color reflected from Bragg lamellae in cells of squid skin. Effective neutralization by anionic screening provides a surrogate for phosphorylation, driving reflectin’s proportional assembly *in vitro*. A physical mechanism for charge regulated assembly of reflectin is proposed.

